# Building the bioeconomy: A targeted assessment approach to identifying biobased technologies, challenges and opportunities in the UK

**DOI:** 10.1101/2023.08.28.554388

**Authors:** Claire Holland, Philip Shapira

## Abstract

We explore opportunities, challenges, and strategies to translate and responsibly scale innovative biobased technologies to build more sustainable bioeconomies. The pandemic and other recent disruptions have increased exposure to issues of resilience and regional imbalance and raised attention to pathways that could shift production and consumption regimes based more on local biobased resources and dispersed production. The paper reviews potential biobased technologies strategies and then identifies promising and feasible options with a focus on the United Kingdom. Initial landscape and bibliometric analyses identified 50 potential existing and emerging potential biobased technologies. These technologies were assessed for their ability to fulfil requirements related to biobased production, national applicability, and economic, societal, and environmental benefits, leading to identification of 18 promising biobased production technologies. Through further analysis and focus group discussion with industrial, governmental, academic, agricultural, and social stakeholders, three technology clusters were identified for targeted assessment, drawing on cellulose-, lignin-, and seaweed-feedstocks. Case studies for each of these clusters were developed, addressing conversations around sustainable management and the use of biomass feedstocks, and associated environmental, social, and economic challenges. These cases are presented with discussion of insights and implications for policy. The approach presented in the paper is put forward as a scalable assessment method which can be useful in prompting, informing, and advancing discussion and deliberation on opportunities and challenges for biobased transformations.

## I. Introduction

Transitioning to bioeconomy systems built on renewable inputs and circular resource streams (EC, 2012) is a multi-level challenge, requiring shifts in production as well as consumption patterns and involving not only high-level policy changes but also meso- and micro-changes in technologies, management systems, and business and governance models. Countries (and regions) will need customisation to their circumstances. In the case of the United Kingdom (UK), as we subsequently discuss, we suggest that there is a realistic vision of future manufacturing that involves supply chains that are substantially less fossil-fuel dependent and more distributed and resilient, relying significantly on biobased and sustainable native resources adapted to local industrial contexts, resources, and biodiversity. Achieving this prospect would be aided by the adoption of innovative biobased technologies for the synthesis and manufacture of chemicals and materials (biomanufacturing), and for applications in agriculture and energy. Yet, while promising, there are significant operational challenges to developing viable and scalable pathways and strategies for biomanufacturing and other biobased production strategies. These challenges include applicability to regional conditions, feasibility, regulatory constraints, and economic-, societal-, and environmental-sustainability aspects, as well as stakeholder acceptance and associated path dependency, institutional, and policy issues.

This paper explores opportunities, challenges, and strategies to translate and responsibly scale innovative biobased technologies to build a sustainable bioeconomy. The European Commission (EC) describes the bioeconomy as the production of renewable biological resources and the conversion of these resources, and their resultant waste streams, into value-added products, such as food, feed, biobased products, and bioenergy (EC, 2012). In the paper, we demonstrate how a rapid targeted assessment method can provide insights about the how the adoption of models based on biotechnologies can address responsible and sustainable bioeconomy development, including anticipation of potential roadblocks to implementation and possible resolutions to overcome these. We proceed by first reviewing the contextual framework for considering bioeconomy development, including environmental, economic and societal aspects, with attention to the UK bioeconomy. This is followed by consideration of the approach and methods used to undertake the targeted assessment, presentation of results, discussion of implications and policy insights, and conclusion and limitations.

## II. Literature Review

### A. Environment, economy, society, and the bioeconomy

The detrimental effects of fossil fuels on the environment and climate change are well documented; CO2 emissions from fossil fuels are reported to contribute to 89% of all emissions (IPCC, 2018). Recognition and societal concern about unsustainability of the continued long term economic dependence on coal, natural gas, and oil has increased (Boehlje & Bröring, 2011; Pfau et al, 2014). Yet, while awareness has risen, adopting sustainable solutions has proven difficult. The piecemeal progress made globally has been further impacted in recent years by the COVID-19 pandemic and rising international conflicts, exacerbating the challenges of achieving targets for net zero greenhouse gas emissions and sustainable development goals (Naidoo & Fisher 2020; Oldekop et al. 2020; Martín-Blanco et al., 2022; OECD 2023).

The reduction of greenhouse gases can be aided by measures such as carbon capture or by changes in mobility and other consumption patterns. However, deeper sustainability transitions necessitate the implementation of rigorous and consistent bioeconomic strategies, including the adoption of innovative biotechnologies (Stark et al., 2022; Losacker et al, 2023). Yet progress here is slow. The present economic size of the bioeconomy depends on definitions used. Estimates using European Union bioeconomy designations indicate that the bioeconomy has maintained a share of about one-tenth of Europe’s (EU-27) economy as reported in 2012 (EC, 2012) through to 2017 (Ronzon et al., 2022). In other words, while there are some country variations, bioeconomy transition is still at an early stage at best. Nonetheless, there continues to be policy ambition to accelerate sustainable growth throughout the bioeconomic value chain – from feedstocks to bioproducts – through the production of renewable biobased resources and the conversion of waste streams into value-added products.

One of the underlying challenges to bioeconomy development is environmental regulation that demands a disproportionately high environmental standard for biobased products compared to their fossil fuel-based counterparts. For example, mandatory ecological sustainability provisions, outlined in both the 2009 EU Renewable Energy Directive (2009/28/EC) (EP, 2009a) and the EU Fuel Quality Directive (2009/30/EC) (EP, 2009b), stipulate that biofuel, produced from biomass, must produce at least 35% less GHG compared to fossil fuels. In 2017, this became at least 50%, and it continues to rise (Priefer et al., 2017) limiting the potential of available biomass and allowing fossil fuel usage to persist uncontested. The evolution of a competitive and sustainable bioeconomy requires significant and enduring public investment in both research and innovation, and the establishment of a pro-active regulatory framework that incentivises private sector investment in the development and commercialisation of new biobased products (Zilberman et al., 2013)

At the same time, it is often assumed that sustainability and “green-ness” are inherent to, and inevitable products of, the bioeconomy (Pfau et al., 2014). Multiple criteria and indicators for a sustainable bioeconomy have been developed (EC, 2015; Fritsche & Iriarte, 2014) but an internationally agreed set of criteria for a sustainable bioeconomy still does not exist. Moreover, the sustainability of bioproduction is not automatically guaranteed. Environmental trade-offs can be required as bioproduction seeks industrial viability (e.g., water usage, habitat destruction, or energy requirements). Environmental repercussions may be heightened if the bioeconomy is developed in isolated ways and in the narrow pursuit of scale and competitive advantage in conditions of growing resource scarcity and demand (Kitchen & Marsden, 2011).

Among other major problems currently faced in bioeconomy development are the geographical availability of suitable bioresources, protection of bioresources from environmental and mechanical damage, reliability of supply, and the complex nature of natural fibres, i.e., differences in consistency, quality, and usability of resources. Although, renewable feedstocks can be produced indefinitely at a certain level, they are still exhaustible if the rate of extraction is faster than the rate of regeneration (Conrad & Clark, 1987). It has been contended that a bioeconomy producing products at a high enough level to fulfil demand could be unsustainable, with fears that this would result in an accelerated rate of depletion of natural resources (water, nitrogen, phosphorus, etc.), land use competition with food production, increased eutrophication, and loss of biodiversity through the exploitation of marginal land (grassland, peatland, wetland, etc.) and forests for agriculture (Langeveld et al., 2010; Landeweerd et al., 2011; Raghu et al., 2011; Sheppard et al., 2011; SDC, 2011; Bringezu et al., 2012; Nuss & Gardner, 2013). We anticipate that such challenges may be more readily overcome by developing supply chains that are more distributed and resilient, relying on biobased and sustainable native resources. For these resources to be sustainably managed and preserve biodiversity, bio-based production ideally needs to be locally managed using viable local resources.

While the bioeconomy has the potential to “green” the economy, the mode of its implementation is paramount to the ultimate success (or failure) on this front. There are two envisaged trajectories for development of the bioeconomy. The first is a delimiting path, involving integration of the bioeconomy into the current economic model where a limited number of actors narrowly define problems and solutions. This would likely produce a bioeconomic model overlain with green claims and where economic output is prioritised over social and environmental impact. A contrasting development strategy, which is the approach that we pursue, is one that is sustainability-centric, incorporating diverse stakeholder and expert input and encouraging multiple solutions (Kershaw et al., 2020). This would require influencing micro-economic behaviour and processes to embed production-consumption chains that are more distributed and resilient (Sonnino & Marsden 2006; Milone & Ventura 2010) as well as the development of complex networks of viable businesses – including small and mid-size enterprises (SMEs), and economic activities that use ecological resources in sustainable, diverse and efficient ways, prioritising a circular economy model.

Such holistic approaches to the bioeconomy view it as a comprehensive social transition that utilises new biological knowledge for commercial and industrial purposes, and to improve human welfare (Enriquez-Cabot, 1998). While social inclusiveness and acceptance are important precursors to the development of a successful bioeconomy strategy, implementation to date has been weak. Bioeconomy sustainability assessment tools do not adequately capture social aspects (Philp & Winickoff, 2018), and social outreach often focusses on creating acceptance for priorities that have already been set rather than inclusive participation and active involvement (Kircher, 2012).

### B. The UK bioeconomy context

The Climate Change Act (UK Parliament, 2008) committed the UK to an 80% reduction in carbon emissions, relative to 1990, by 2050. In June 2019, this target was extended to a 100% reduction. The UK is not on track to meet either of these targets. Meanwhile, the UK’s “Bioeconomy Strategy: 2018-2030” aimed to double the impact of the UK bioeconomy from £220 billion (2014) to £440 billion by 2030 (BEIS, 2018).

Subsequently, the global COVID-19 pandemic (early 2020 through to May 2023) disrupted many plans, with contrasting implications. On the one hand, the pandemic led to the biggest economic crisis in the UK since records began, with a fall in GDP of 20.4% in the second quarter of 2020 (ONS, 2020). Economic problems persisted into 2021, although with some recovery towards the latter part of the year, such that by November 2021 the UK economy had regained its pre-pandemic size (Keep, 2022), albeit with variations by sectors and regions and with high levels of ongoing uncertainty (complicated not only by the pandemic but also by Brexit, rising fossil fuel prices, and global disruptions). On the other hand, the pandemic intensified debate, and led to government policy announcements, about “building back better” with growth that levels regional imbalances, fosters resilience, and transitions to net zero greenhouse gas emissions (HM Treasury, 2021). Such policies also featured in the 2021 UK Innovation Strategy (BEIS, 2021), which superseded the 2018 UK Bioeconomy Strategy, highlighting the potential of engineering biology as an enabling technology in rebuilding the economy.

The heightened debate about more sustainable and resilient production and consumption transitions in the UK is a significant development. Relying on requests for behavioural change by itself will be insufficient and will not enable the UK to resist future crises. Instead, structural changes in industry and the economy, solid commitment to environmental targets, and a levelling of societal imbalances are needed. In considering how a greener, redistributed recovery and transition might occur, possibilities have been raised that involve shifts to production regimes based more on local bioeconomy resources and dispersed production, and less on point-source resources (fossil fuels) and distant international sources, with customisation to local contexts, industry, and biodiversity. Innovative biological pathways for making chemicals, materials, and other bioproducts, and for applications in agriculture and energy have emerged that promise not only greater sustainability but also new functionalities. However, any shift to biobased is a multi-dimensional challenge, with issues that are not just technical and environmental but also political, economic and societal.

A key aim of this paper is to go beyond high-level pronouncements and broad statements of intent to examine more specifically which transitions to sustainable biobased production are most promising, how these transitions might occur, what challenges might arise, and how these challenges could be overcome. We take these up through developing and implementing a targeted assessment method applied to potential UK biobased production pathways.

## III. Methods

There are multiple technology assessment approaches to evaluating potential technological developments and impacts and consider policy implications, including approaches relying on extensive expert analyses and those which involve stakeholders and societal actors (Matthews et al., 2019). We pursued a simplified and low-cost approach that combined analysis of available documentation, iterative feedback, and focus group stakeholder participation. The assessment approach was targeted in that, after an initial broad scan of potential bioeconomy technologies, it concentrated analysis on a small number of selected cases of biobased production technologies. The steps in the method are described in this section and presented in Figure 1. Further details are included in the supplementary materials, including search terms and selection processes. This targeted assessment method was implemented over a period from August 2020 (start of long-list generation) to April 2021 (completion of case studies).

**Figure 1.**
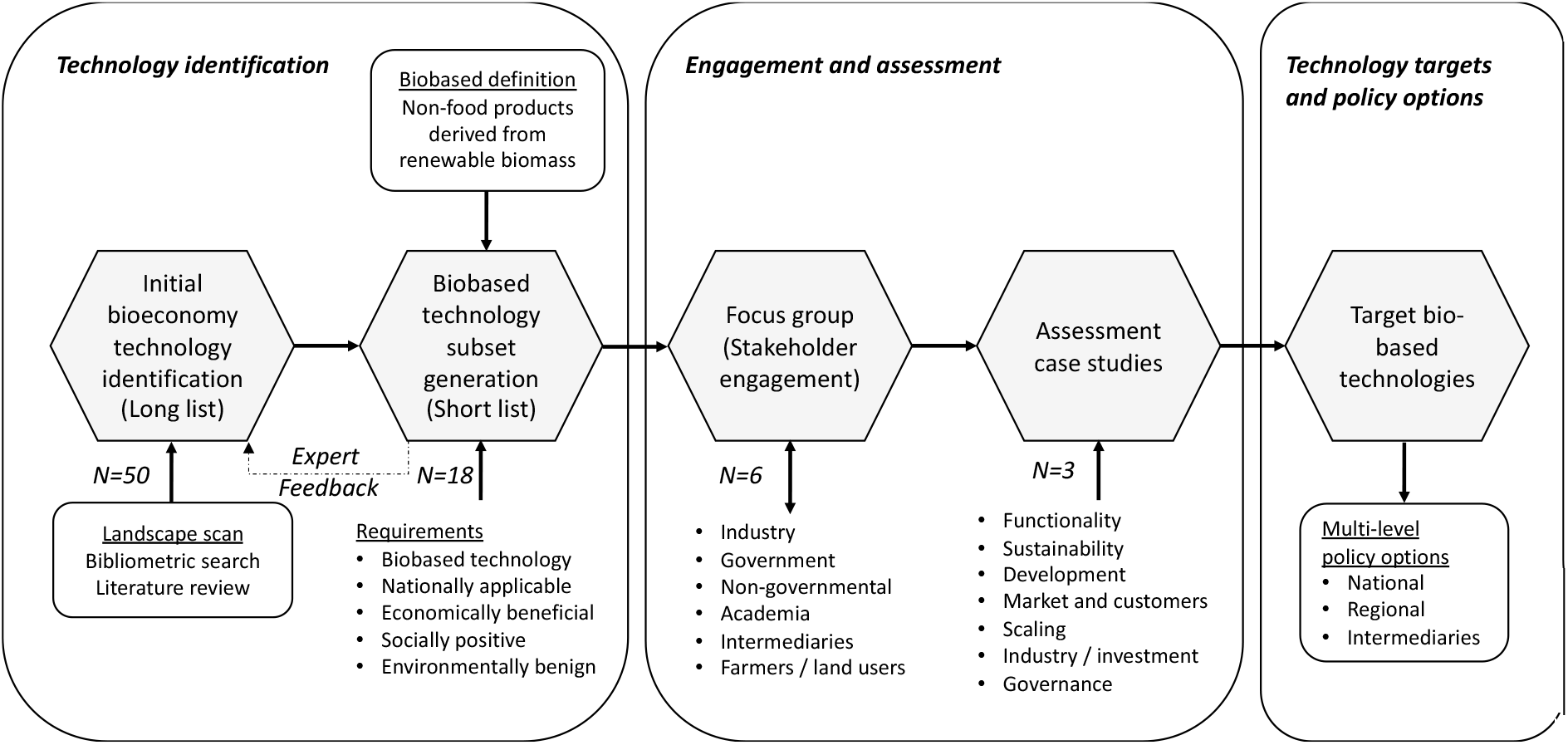
Targeted Assessment Approach.

### A. Assessment steps

#### 1) Bioeconomy to biobased production technologies: long list to short list generation

Through landscape analysis and bibliometric searches, we developed a starting long list of extant and emerging innovative technologies that could contribute to bioeconomy development. Bioeconomy technologies were broadly defined as ones that facilitated the production of goods, materials, energy, and services with the sustainable use of biological resources. We used a combination of bibliometric searches, reviews of available literature, and searches of other online resources. There was a focus on technologies that were generic (i.e., not highly company specific) and either entering or relatively close to market – i.e., which could be implemented within, as a maximum, the next 5-10 years. A total of 50 potential bioeconomy technologies emerged (see Supplementary Materials and Table S1).

To refine this initial bioeconomy set to a short list of promising biobased production technologies, we applied a definition of biobased as “non-food products derived from renewable biomass (plants, algae, crops, trees, marine organisms, and biological waste from households, animals and food production)” (ERRMA, 2007). Given the saturation of the UK market with biobased technologies for healthcare applications, only technologies pertinent to bioagricultural and bioindustrial applications were included in this study. Long list technologies were further assessed for their ability to meet requirements related to national applicability (in this case, to the UK). Criteria to assess the promise of the technology included utilisation of native feedstock, resources, and expertise and the potential to be economically beneficial (either by reducing dependence on imports, bolstering a particular market, or providing new goods or functionalities), socially positive (including allowing local management of resources, job creation, or other societal gains) and environmentally benign (or the potential to be so) considering such aspects as greenhouse gas emissions, toxicity, resource depletion, habitat destruction, and water use.

As the list of candidate technologies was generated by the author research term, it was presented to experts in biotechnology, engineering biology, and science and technology policy (four workshops or seminars). The feedback received contributed iteratively to refinements of initial lists, with 18 technologies determined to best fulfil the requirements criteria and included on the short-list of promising biobased production technologies. (See Results section and Supplementary Materials).

#### 2) Focus group (stakeholder engagement)

As part of the assessment approach, a focus group process was initiated to engage with stakeholders. Exemplary technologies were selected for further discussion and feedback in the focus group. The focus group facilitated a multi-discipline discussion about the feasibility, implications, and considerations associated with each exemplary technology, and identified associated roadblocks and considerations relating to its development and to its contribution and integration into the UK bioeconomy. The focus group (held online in January 2021) was attended by academic, scientific, governmental, regulatory, think-tank, policy, agricultural, and industrial stakeholders (10 attendees plus 2 research team members). Participants were asked to address three principal questions: (1) What are the specific challenges you see relevant to each of these specific technologies? E.g., technology, management and/or regulation. (2) What are the main governance, funding and regulatory roadblocks you consider to be the most important and how can these be overcome? (3) Are there any other technologies you think should be included as exemplary case studies based on the UK’s native feedstocks, resources, expertise or policy context?

#### 3) Assessment case studies and target biobased technologies

Following feedback from the focus group, three technologies were highlighted as presenting the most promising options for biobased production. The focus on three technologies was intended to allow for the development of in-depth case studies to pilot a targeted assessment method that could be made available for subsequent scale up to other biobased technologies. The case studies were undertaken using secondary data sources with a focus on the following probes: (1) Functionality: Technical specification, novel functions, and comparison to the status quo. (2) Sustainability: Source of feedstock, ability of UK to provide / import, and sustainability/environmental issues of production and consumption. (3) Development: Current use, and alternative and developing methods of utilisation, extraction, and valorisation. (4) Market and customers: Affected markets and customers, current and potential scale / growth / locations of the market, and potential market competitors. (5) Scaling: Potential for and requirements of scalability of the extraction/ valorisation process, and examples of technologies in development. (6) Industry, investment and partnerships: Current industry structure, potential for redistributed (small-scale, decentralised) production, and capital requirements. (7) Governance: Regulatory aspects, public acceptance and possible opposition to this feedstock / technology. As part of each case, for each of the pilot target biobased technologies, consideration was given to multi-level policy options to foster responsible and sustainable development.

## IV. Results

### A. Long list to short list

As noted, a long list of 50 extant and emerging innovative bioeconomy technologies was identified through bibliometric and documentary analysis and expert workshop or seminar feedback. Review of these emerging and innovative biobased technologies yielded company-specific products, specific emerging technologies at varying development stages, and pathways to generate already biobased end-products. Technologies that did not fulfil the definition of biobased were removed, including solar, wind, and alternative energy-based technologies.

Of the initial 50 biobased technologies, 33 were progressed further. These technologies were assessed (drawing on expert input) for their ability to fulfil the five requirements for subset generation (see section III.A.1). Although some technologies are not presently environmentally benign, they were assessed on their capacity to be so with further development to balance the immediate need for these technologies with long-term potential. Eighteen technologies fulfilled the requirements criteria, and from these six exemplary technologies were selected for further discussion (biomethane, high purity lignin, MFC/ nanocellulose, PHAs from fatty acids and urban waste, seaweed technology, and bioethanol) via a focus group. While not necessarily the most innovative of the technologies available, they were selected to represent a range of different feedstocks, low- and high-value products, and a diverse range of primary end uses, from energy, to biomaterials, to platform technologies.

### B. Focus group findings

In the focus group discussions, an early consensus was established that subsequent case studies should focus on specific native feedstocks, with feedstock-specific technologies considered within this overarching context, to address the conversation around management and utilisation to ensure sustainability. The downstream uses/technologies of each feedstock platform are extremely diverse, as are the environmental, social, and economic impacts. Consideration of environmental, social, and economic challenges, the potential for scale-up of a specific resource, and optimisation at the primary stage can influence products downstream but not vice versa. As such, a ground-up approach became the focus.

A core challenge envisaged for biobased feedstocks is that of supply chain, which needs to be sufficiently large to accommodate commercial demand whilst having a definitive economic and environmental impact. Due the size of the UK and resource availability, it was recognised that it would be relevant to focus on small-scale, high-value opportunities. However, for there to be a large-scale effect on climate change, the UK would also need to export technologies where the balance is reversed (i.e., large-scale, low value). Utilisation of waste streams may be a way to negate this issue.

Activation barriers were identified as a potential roadblock. Extant systems with integrated supply chains, such as those reliant on fossil fuel feedstock, can function at a given economic scale. For these systems to transition to biobased feedstock, different supply chains may need to be established, potentially at a different scale, and requiring investment in alternative infrastructure, expertise and facilities. Such a transition would require both Government promotion, through initiatives and incentives for companies to develop and implement more sustainable biobased technologies, and societal pressure to encourage companies to expedite this shift. A supportive and consistent government policy – such as carbon pricing/ the voluntary carbon standard – is needed with a framework that supports the transition to a low-carbon world.

### C. Case studies

Based on focus group feedback, assessment case studies were developed for cellulose, lignin, and seaweed to assess in greater detail the potential and scope of these feedstocks as the foundation for development of a sustainable biobased UK. The case studies address the seven categories indicated in section III.A.3 (see also Supplementary Appendix).

#### 1) Functionality

Cellulose – the most abundant natural biopolymer with ∼100 billion tons/annum available globally – is predominately derived from plant cell walls but is also present in many algae and oomycetes and is secreted by some bacteria. Cellulose is endowed with numerous attractive and exploitable properties – it is strong, insoluble in water and the majority of organic solvents, and biodegradable (Hepworth & Bruce, 2000). The usability and properties of cellulose depend on its degree of polymerisation, with long-chain and short-chains being preferable for different end-products.

Lignin, another plant-derived polymer, is the second most abundant natural polymer on Earth with ∼ 20-billion tons/annum available globally (Fernandez-Rodriguez et al., 2017). Lignin has potential as a low-cost carbon precursor (Rosas et al., 2014) as it comprises 30% of all non-fossil organic carbon, is CO2 neutral, antimicrobial and biodegradable (Boerjan et al., 2003), and is the only known renewable source of extractable aromatic compounds from biomass (Boudet et al., 2003). Cellulose is commonly complexed with lignin, and hemicellulose as part of lignocellulosic biomass. Valorisation of this lignocellulosic biomass has numerous benefits; allowing the production of added-value chemicals, materials and fuels from a global renewable resource, thereby decreasing dependence on petroleum, and simultaneously improving the economics of existing lignocellulosic processes in the biofuel, and pulp and paper industries (Chen and Wan, 2017; Liu et al., 2018).

Seaweed is of interest for applications in food, chemicals, cosmeceuticals, nutraceuticals, bioenergy and biofuels, due to an increased interest in “sea vegetables”, variety of cultivars, global bioresource availability, and the development of valorisation processes and higher-value biobased products (Cefas, 2016). While similar in many respects to terrestrial plants, there are differences in seaweed composition and development. For example, macroalgae have a higher concentration of many minerals and nutrients – including Ca, K, Mg, Na, Cu, Fe, I and Zn – compared to terrestrial plants in concentrations up to ten-times higher (Mouritsen, 2013). Seaweeds also typically have faster growth rates, higher productivity, and higher polysaccharide contents – similar in structure to that of land plants but different in abundance ratios.

#### 2) Sustainability

Extraction of lignin and cellulose is possible from first- and second-generation biomass, waste, and as a by-product of the pulp and paper industries and bioethanol production. The paper and pulp industry is the main producer of commercial grade lignin, with Kraft pulping alone producing ∼130-140 million tons of lignin per annum (Rinaldi et al., 2016; van den Bosch et al., 2018). However, most of this is burnt to produce on-site heat/energy; only ∼2% is used for higher-value applications (Thielemans et al, 2002; de Wild et al, 2014). Recycled paper is another viable source of lignocellulose; 5.5 million tonnes of paper and cardboard were used in the UK of which 65.6% was recycled (DEFRA, 2021). Paper fibres undergo recycling a limited number of times before fibre quality is compromised, so annual additional waste-paper residues amount to ∼1.16 million tonnes (NNFCC, 2014). The UK also exports ∼4.2 million tons of paper for processing, which could instead be utilised (NNFCC, 2014). Another source of cellulose is bacteria, which produce purer cellulose that has a higher water content, and higher tensile strength (due to a higher degree of polymerisation) (Klemm et al., 2005) than plant-derived cellulose. This form of cellulose can be produced from numerous types of bacteria, usually via fermentation or reactor-based production, although this is not currently possible on an industrial scale.

Woodland is an important source of lignocellulosic biomass, but also for carbon sequestration, biodiversity, and as a natural system for flood and soil-loss prevention. Total UK woodland is estimated to be ∼3.21 million hectares (Forestry Commission, 2020). It has been determined that the proportion of woodland residues that can be removed without affecting soil stability and nutrient levels is around 50% (Searle & Mallins, 2013). Meanwhile, agricultural land accounts for around 62.8% of total land use in the UK (MHCL, 2020). First-generation crops and food-feedstocks are unsuitable for large scale production of bioproducts as: (1) they typically require good agricultural land and fertile soils, which is limited (Rosengrant et al., 2013; Pedroli et al., 2013), and (2) competition for use can contribute to food scarcity, and inflated food and animal feed prices (Sims et al, 2010). Despite concerns about food security, the Waste Resource Action Programme (WRAP) estimates ∼3.6 million tonnes of surplus food and food waste is generated annually on UK farms alone, equating to around 7.2% of total UK production, with an economic value of ∼£1.2 billion. This could be used as a source of lignocellulose (WRAP, 2019). Interestingly, seaweed can help reduce emissions from agricultural, which is responsible for around 30% of all GHG emissions resulting from land use (Duarte et al., 2017), by reducing dependence on environmentally damaging synthetic fertilisers (Smith, 2002), improving crop productivity and soil amelioration (Roberts et al., 2015; Smith, 2002), and through use as a prebiotic food additive to reduce methane emissions from cattle (Robertson et al, 2000; Smith 2002), enhance livestock health (Rey-Crespo et al., 2014; Makkar et al 2016) and substitute antibiotics (O’Doherty et al 2010, O’Shea, 2014).

The UK produces nearly 16 million tons of biomass waste per year, predominately in the form of municipal green waste (MGW). MGW is an abundant and consistent (both seasonally and spatially) source of lignocellulosic material (estimated content: 35-50% cellulose, 31-42% hemicellulose, 3-35% lignin). Nearly 5 million tons/year of green waste are collected in the UK, accounting for 31% of the total waste biomass generated per annum (NNFCC, 2014). Waste availability and sustainability depends on human consumption practices, availability of natural resources, and the diversion/processing of viable lignocellulose sources away from landfills (Clark and Deswarte, 2008), reducing the detrimental environmental, economic and social impacts associated with them (Sharholy et al., 2008; Venkata Mohan S., 2016; Nizami et al., 2016b).

Seaweed is not compounded by the same land use issues as lignin and cellulose. As a viable source of both food and third-generation biomass, seaweed confers numerous environmental advantages, including acting as an anthropogenic CO2 sink, regulating ocean pH and natural oxygenation and nutrient cycles, waste purification (FAO, 2014), bioremediation, and promotion of biodiversity (Buschmann et al., 2017; Duarte et al., 2017; Vásquz et al., 2014; Steneck et al., 2002; Smale et al., 2013). In addition, farmed seaweed is considered a carbon negative crop (Duarte et al., 2017). Problematically, assessment of seaweed sustainability in the UK is compounded by a lack of information about the potential effects of scaling in European waters, invasion of non-native species, the risk of pathogens, losses to other industrial sectors (such as leisure and fishing), and competition with ecosystems that provide vital carbon sequestration services, such as seagrass meadows (Duarte et al., 2005, 2013; McLeod et al., 2011; Fourqurean et al., 2012). Proper evaluation and assessment of suitable locations for farms or harvesting in the UK is needed so potential conflicts can be mitigated and opportunities for development can be exploited. Data pertinent to productivity, yields, and optimal algal cultivars is needed to ascertain the scale of UK production that can be sustainably harvested.

#### 3) Development

Lignin, cellulose, and seaweed can all be considered bulk feedstocks. While they can be utilised as raw materials, it is through degradation to their component parts, and purification of these moieties, that they can be utilised for higher-value products. For example, for effective valorisation via fermentation or biotransformation, lignin must be broken down to mono-or oligomers to allow utilisation of its carbon. Different depolymerisation methods yield different/ higher fractions of specific phenolic monomers, including vanillin, guaiacol, p-coumarate, benzoate and catechol, which are of particular relevance for downstream biochemical conversion processes. As such, the depolymerisation method needs to be optimised for the desired end-product, with preference given to methods that retain the integrity of other components to allow maximum yield of usable material from the original feedstock. The most effective pre-treatment methods for lignocellulosic material – physical and chemical – often produce unwanted by-products, causing environmental damage or inhibiting hydrolytic enzymes downstream. By contrast, biological pre-treatments are essentially environmentally impact free: exploiting enzymes or accessory proteins. Enzymatic extraction methods are in development, with numerous potential candidates for biochemical and synthetic biology approaches, including hydrolytic and glycosidic enzymes, cellulases, and cellulose-associated proteins (Lui et al., 2015; Holland et al., 2019).

The UK has a long history of the harvesting of wild seaweed, but seaweed farming is not currently a well-developed industry. There is significant potential for the development of the seaweed sector given the UK’s extensive coastline and expertise in macroalgal research, including algal taxonomy, cultivation, physiology, metabolism, biochemistry, and molecular biology (Cefas, 2016). However, development of the UK seaweed sector is limited by numerous problems, including how to establish the industry at a large-scale sustainably, economic viability (especially considering the low cost of imports from Asia), and development of high-value products. For example, expansion of seaweed aquaculture into offshore areas – allowing greater volumes to be cultivated – raises challenges in engineering technology able to withstand harsh environments and high salinity, as well as issues of decentralised energy supplies, reduced nutrient accessibility, and additional costs for transport and a skilled workforce. Aspects such as storage, pre-treatments, a viable supply chain, and the use of algal bioreactors for cultivation need to be developed further (Cefas, 2016).

A particularly interesting area of development, which has no counterpart in the fossil fuel industry, is the use of synthetic biology and genetic modification to impart advantageous properties, or novel characteristics into these feedstocks. Future bioengineering approaches could, for example, impart genetic modifications that either reduce lignin content, allowing greater saccharification of biomass without compromising plant yield or growth, or modify the lignin structure to make it more amenable to chemical or enzymatic degradation under milder conditions (Rinaldi et al., 2016; Ralph, 2010) by replacing recalcitrant ether bonds with more labile ester bonds. Synthetic biology approaches could be useful in the industrial application of these feedstocks, using native and engineered microbes to convert building blocks into bulk chemicals (typically through fermentation), and via biotransformation into smart biomaterials and high-value products (Becker & Wittmann, 2019). For increased industrial application, breeding of efficient microbial cell factories needs to be expanded from the lab- and pilot-scale (Kohlstedt et al., 2018), and microbial strains need to be produced that allow operation at high levels (Overhage et al., 2003; Sonoki et al., 2014), have extended substrate specificities (van den Bosch et al., 2018; Buschke et al., 2011; Sonoki et al., 2014; Vardon et al., 2015), and allow integration of various production pathways.

#### 4) Market and Customers

In 2019 the global lignin market size was estimated at US$954.5 million with an expected CAGR of 2.0%, in the 2020 to 2027 period (Becker & Wittmann, 2019). Demand for kraft-derived lignin (ligno-sulphonates) currently accounts for the largest market share of total lignin, accounting for 85.5% of demand in the US in 2019, with an expected CAGR of 1.6% between 2020 to 2027 (Becker & Wittmann, 2019). The global cellulose market is significantly smaller, valued US$219.53 billion in 2018 (FBI, 2020) and US$211.68 billion in 2019 (GMI, 2020). It is projected to rise to US$305.08 billion by 2026 with a CAGR of between 2.9-4.2% while, the seaweed cultivation market had an estimated value of US$16.7 billion in 2020, with a CAGR of 12.6% (Ferdouse et al., 2018). Global production of marine macroalgae, both wild-collected and cultivated, in 2018 was reported to be 32.4 million tonnes, of which 97.1% was from seaweed farms (FAO, 2020).

At present, markets for bioderived feedstocks can be broadly categorised into three groups: (1) direct use, (2) drop-in replacements for oil-based products, and (3) as a raw material for commodity or high-value chemicals, and value-added products. Lignin feedstock is predominately used for animal feed, and in the production of macromolecules used in biofuels and biorefinery catalysts (Becker & Wittmann, 2019) but, it can be used directly or for the production of a wide variety of chemicals and materials. Products from delignification can be used directly (e.g., concrete/cement additives, glue, absorber and fertilisers; use depends on the delignification method), or can undergo pre-treatment and depolymerisation, via pyrolysis, cracking, solvolysis, acid/base catalysis, enzymatic or hydrogenolysis methods. Alternatively, these oligomers can go through aromatic funnelling to produce high value aromatics such as catechol and protocatechuate (Vardon et al., 2015; Wu et al., 2017). Further processing is possible via biotransformation or fermentation, leading to direct use products for chemicals, fuels, materials and flavours.

Cellulose is a major constituent of both the paper and textile industries, but the development of novel cellulose-based products has expanded its market potential. Regenerated cellulose can be used for the manufacture of rayon, cellophane and viscose, textiles, medical devices, nitrocellulose, and artificial membranes. Microcrystalline cellulose is used in the consumables, food, and pharmaceutical markets as E-numbers, inactive fillers, and as emulsifiers, thickeners and stabilisers. Other potential uses of cellulose include as piezoelectric sensors, housing insulation and fireproofing, water-soluble adhesives and binders, filters and membranes, bulking agents for cement, as biodegradable matrices for the preparation of nanocomposites, and as reinforcing fillers for polymer-based nanostructured composites, thermoplastics and thermosets.

The lack of a diversified market for seaweed, and access to cheaper raw materials and labour from Asia, means there is significant competition and lack of opportunities/incentives for seaweed producers in Europe (Cefas, 2016). This could be helped by opening new markets, valorisation of seaweed to higher value and specialty products, and a substantial increase in seaweed production, which could be achieved by near-shore farming in the short-term, and further off-shore in the long-term as technology is developed to support this. It should also be noted that the monetary value of non-market public goods, such as ecosystem services, bioremediation, and biodiversity, are difficult to quantify and vary significantly dependent of seaweed site, species, and cultivation method (Cabral et al, 2016).

#### 5) Scaling

The upscaling and commercialisation of biobased technologies face several issues: availability of sufficient feedstock, the chemical recalcitrance and structural complexity of the feedstock, purity, the type and quality the feedstock, and biocompatibility. This is true of lignin, cellulose, and seaweed. For sustainability and economic viability, upscaling needs to focus on biobased feedstock from extant side streams and waste, and development of cascading routes of production. Cascading routes can be used to prioritise production of high-value products, with waste and by-products of this process being valorised downstream to produce lower-value or bulk products. Using this approach, products can be removed and used at various stages of the production process depending on desired end-use or physical, chemical, and mechanical properties. This cascading process requires optimisation of both upstream (e.g., delignification bioengineering, extraction) and downstream (e.g., depolymerisation, scale-up, cascading) processes (Rinaldi et al., 2016) taking into account environmental, social and economic aims. Both “high volume, low value” and “low volume, high value” applications are needed for the full economic potential of these feedstocks to be realised (Chem. Eng. News, 1984; Strassberger et al., 2014). This allows maximum yield of marketable products for minimum feedstock.

Although pilot facilities for seaweed cultivation have been established in the UK, knowledge about the ecological impacts, yield, and operational costs are not sufficiently robust for real insight into the cost, both economically and environmentally, of scale-up and development. The lack of recent and accurate estimates of the size and composition of the UK’s standing stock is a major problem for development and upscaling due to uncertainty about whether there are sufficient resources for industrial scale application. In 2013 UK seaweed production from wild harvest was estimated at approximately 2000-3000 dry tonnes (AB-SIG, 2013). However, this figure does not include the unknown quantities of storm-cast seaweed and subtidal kelp that are also collected. At present seaweed biomass represents only a small quantity (30 x 106 tonnes; fresh weight) of the total global biomass supply compared to terrestrial crops, grasses, and forests (16x1011 tonnes; fresh weight) (Buschmann et al., 2017). Expansion and scale-up of seaweed aquaculture, especially into offshore areas, would require development of suitable technology and equipment, training of a skilled labour force, and new seaweed cultivators (Duarte et al., 2017). As scale-up of seaweed production may cause a decrease in market prices, discouraging farmers and investors, therefore, growth in seaweed cultivation needs to be coupled with the development of novel applications and high-value products (Mazarrasa et al., 2014).

#### 6) Industry, investment and partnerships

Seaweed aquaculture is currently limited in the UK and existing pilot farms – for example, Queen’s University (Northern Ireland), SAMS and the University of Highlands and Islands (Scotland) and Swansea University (Wales) – mostly remain in the research and development stage. There are a number of UK algal projects (e.g., EnALgae, At∼Sea, BioMara, SeaGas, MacroFuel, MacroBioCrude) with aims to produce biobased products from algae, while Wave Crookes has become the first commercial UK sea farm. Around 80% of global commercial macroalgae is used for human consumption (West et al., 2016) but a wide range of products can be obtained from seaweed, both direct use, including human and animal feed, chemicals (e.g., hydrocolloids, phytocolloids, carrageenan, agar), fertilisers, bioactives for nutraceuticals and dietary supplements, cosmeceuticals, food ingredients (such as omega-3 fatty acids), natural food colourants and dyes, fertiliser, bioplastics, specialty chemicals (such as phycocolloids), and chemical feedstock, pharmaceuticals, probiotics (Gomez-Zavaglia et al., 2019), biofuel, and for bioremediation and purification services. Despite the opportunities for seaweed as a feedstock, it is not without its challenges, significant research and investment needs to be made into the development and optimisation of the industry, including selection of preferential species for specific end uses, and for sustainability and economic considerations (Zhang et al., 2007; Robinson et al., 2013; Li et al., 2016). Aspects such as storage, pre-treatments, a viable supply chain, and the use of algal bioreactors for cultivation need to be investigated further (Cefas, 2016).

Most commercial technical lignin is produced by MeadWestvaco Cooperation and Borregaard Industries in the form of kraft pulp. While there are no significant, independent, UK companies creating technical lignin, the chemical industry is well established in the UK and integration of lignocellulosic feedstock into existing production streams is feasible. While the products of kraft pulping (mostly sulfonated) can be used directly as “low-value, high volume” additives for concrete, glue, absorbers, emulsions, and fertilisers (Cao et al., 2018; Khan et al., 2018; Naseem et al., 2016; Schutyser et al., 2018) the majority is burnt for energy generation rather than undergoing valorisation as they are unsuitable for many high-value applications (Pye, 2010; Schutyser et al., 2018). The need for technical lignin suitable for valorisation to higher value products (such as vanillin, phenol derivatives, phenolic resins, and BTX), and the notoriety of the Kraft process as environmentally damaging – it uses harsh chemicals, large quantities of water, and emits organosulphur compounds (Ragnar et al., 2014; Ek et al., 2009) – could boost demand for Organosolv-derived lignin. The Organosolv method is both more environmentally benign and produces higher yields of high-purity lignin. Despite the efficacy of the Organosolv method, which results in a high degree of lignin removal (>70%) with minimal loss of cellulose (<2%) (Rinaldi et al., 2016), industrial Organosolv pulping enterprises have mostly been limited to the pilot scale, with few lasting longer than 5-years. A notable exception is the Alcell process developed by Repap Enterprises Inc (Rinaldi et al., 2016). The inability of Organosolv to saturate the market, despite its competitive isolation ability and lower environmental toll, is thought to be due to two main factors: (1) establishment of Kraft mills are a multi-billion pound investment, meaning further investment and regulations would be needed to both fund and incentivise a change in the industry (Rinaldi et al., 2016), and (2) a lack of high-value applications able to utilise the high-quality, sulphur-free lignin produced (Michels & Wagemann, 2010; Viell et al., 2013).

Cellulose derivatives include, among others, commodity-use cellulose pulp, fibres, ethers, esters, microcrystalline cellulose and nanocellulose, and regenerated cellulose, which are used in the textiles, food, chemical, pharmaceutical, construction, paper, pulp, and materials industries (FBI, 2020). There are a number of UK business established across these sectors, notably Cellucomp, FiberLean Technologies (Imerys and Omya), and Zelfa Technology. Cellulose for industrial use is mainly obtained from wood pulp (via the kraft process) and from cotton (Klemm et al., 2005). Cellulose derivatives hold three core advantages for the industry over their oil-based counterparts; cellulose as a feedstock is bio renewable, widely available from multiple sources, and comparatively low cost. However, there are disadvantages to cellulose and natural fibres that limit their industrial application: variable quality, limited maximum processing temperature, incompatibility with hydrophobic polymer matrices, aggregate formation, lower durability, and poor fire resistance.

#### 7) Governance

A shift to a biobased economy utilising lignocellulosic feedstock will inevitably increase demand on agriculture and woodlands, and it is important that this is done sustainably, which highlights the need for policy direction and incentives tied to soil carbon/soil microbiome/plant-soil interactions/soil additives to encourage a greater yield of biomass, carbon sequestration, and protection of biodiversity and limited resources. Human consumption practices need to become more sustainable and effective waste management strategies need to be implemented to ensure the availability of natural resources, and the diversion of viable waste feedstocks away from landfills (Clark and Deswarte 2008), thereby reducing the detrimental environmental, economic and social impacts associated with them (Sharholy et al., 2008; Nizami et al., 2016b).

Significant research and investment is needed to develop and optimise the seaweed industry. The absence of specific regulations for seaweed farming in the UK, which creates uncertainty for investors, alongside the lack of diverse and high value markets for end products, promotion and marketing, means there is insufficient interest from investors to supply the start-up capital needed. Investment is needed for scale-up, development of supply chains, downstream processing, innovation (e.g., strain development, algal bioreactors, seasonality), and infrastructure (e.g., storage, distribution). Economic incentives – such as those linked to climate change mitigation, and bioremediation – and additional funding for seaweed-based research is needed from the Government to incentivise investors and farmers, helping the seaweed industry overcome this hurdle.

## V. Discussion

We have presented a targeted assessment process where, for a single national economy (UK), we identified promising biobased technologies from a larger set of bioeconomy technologies. We evaluated these promising possibilities using assessment criteria related to environmental, economic, and societal sustainability, and discussed them in a focus group. This led to the identification and assessment of three native feedstocks – cellulose, lignin, and seaweed – that could be utilised in the UK to advance biobased economic transition, with scope for differing functionalities and the potential for multifaceted applications.

A core roadblock in the transition towards a bioeconomy is land use and whether there is sufficient biomass available to meet increasing demands. However, as we have discussed, land use issues can be addressed through the development of management practices that prioritise sustainability rather than intensification, and utilisation and integration of core feedstocks from a range of different sources. Rather than utilising biobased feedstock from first-generation biomass, which competes with crops for the food and feed industries, extraction and derivatisation of biomass from second-, third- and fourth-generation biomass should be prioritised. This includes, but is not limited to, waste (forestry, agricultural, industrial biomass-based and municipal solid organic), algae, fermentation of microorganisms, and through synthetic pathways. Waste-based feedstocks are a particularly promising source of lignocellulosic material as they are usually residues for which there is either no commercial demand, or the alternative to valorisation is disposal/ low-value energy generation. Utilisation of waste streams as feedstock rather than higher value commodities can also confer social benefits when the development and management of feedstock is controlled in rural areas on a local scale rather than via nationwide or global management (Kitchen & Marsden, 2011).

At present, we find that the cultivation, isolation, and utilisation of lignin, cellulose, and seaweed feedstocks in the UK is not entirely environmentally benign, although (unlike fossil fuels) they have the capacity to be so with further development and optimisation, and through the adoption of circular economy models, sustainable management, and technological advancement. Our approach here balances the need for immediate change, with long-term potential for development opportunities to further “green” processes, improve economic competitiveness, and production of biobased products and technologies favourable or novel applications/ functionalities. Development of these feedstocks presents a multi-faceted challenge. An obstacle to a biobased economy is the high associated costs, compared to fossil fuel-based alternatives. Although the costs of raw materials are relatively low, the production of biobased products tends to be dominated by subsequent processing costs, reducing their economic competitiveness with entrenched industries and products. By contrast, despite having high raw material costs and initial investment, the fossil fuel industry has low variable and production costs (Dale, 2003; Zilberman et al., 2013) due to sustained research and development over many years. While these variable costs may increase over time, as the finite fossil fuel resources are depleted, this is likely to be too late to avoid the environmental damage predicted for their continued use.

Despite broad scientific agreement on global environmental changes, and the need for global political consensus to engage with sustainable development, the political and regulatory frameworks required to implement the necessary changes have been compounded by a lack of impetus and direction, and entrenched supply chains resilient to change. For all three feedstocks presented here, there is a clear need for increased investment in R&D, more policy incentives to make the bioeconomy more competitive, and for a reduction in fossil fuel subsidies to level-the-field (Levidow et al., 2012; Popescu, 2014; Philp, 2018; EC, 2018).

Research funding mechanisms that explicitly provide for collaboration between scientists and process engineers will help ensure that research is focussed on those areas most likely to significantly reduce processing costs. The main objectives are a close alignment of policy, science, and patents. Public-private partnerships and the creation of technological clusters are given high priority. While funding is strongly focussed on the life sciences, little attention is paid to other areas of research, such as agro-ecological faming, breeding of alternative crops for marginal sites, or social innovations (De Besi & McCormick, 2015). For the three identified feedstocks, support is needed for the establishment and integration of biorefineries, which will allow optimal utilisation and valorisation of feedstock, by-products, and waste streams, thereby yielding maximum product from minimum feedstock.

Technological advances often come from academia but are unable to bridge the gap to become marketable innovations. More focus is needed in the UK to progress discovery science at early technology readiness levels (TRLs 1-2) to closer-to-market technology readiness levels (TRLs 7-9). De-risking of novel technologies is needed with more financial and operational support for start-ups and SMEs, for longer development periods. Investment is also needed for scale-up, development of supply chains, downstream processing, innovation (e.g., strain development, algal bioreactors, seasonality), and infrastructure (e.g., storage, distribution). Even within the circular economy context, economic value is given precedence over other forms of value, such as environmental sustainability, amenity value, and ecosystem services (Nizami et al., 2017b; Stegmann et al., 2020; D’Amato et al., 2020). Economic incentives linked to climate change mitigation, and bioremediation could encourage investors to overcome this hurdle (Cabral et al, 2016).

The assessment of the three feedstocks and their potential to input into bioproduction further surfaced needs for development of policy and legislation that supports the evolution of the bioeconomy in a sustainable way, prioritising environmental and social benefit. This requires a multifaceted approach that both bolsters biobased technologies, for example through support for new markets for biobased technologies, and recognises benefit in excess of marketable goods, such as incentives linked to bioremediation and environmental services, and social and health impacts. Regulation that would, for example, allow crop modifications that produce more lignin, or make lignin more amenable to downstream processing, could be considered. Policy is also needed that levels the playing field with the fossil fuel industry, through initiatives such as carbon taxation, removal of oil subsidies, and legislation to make drop-in replacements more economically viable. A commitment is needed to long term systematic change, rather than short term fixes.

## VI. Conclusions

We have presented a targeted assessment process where, for a single national economy (UK), we identified promising biobased technologies from a larger set of bioeconomy technologies. From a longer list of potential technologies, we identified and evaluated promising possibilities using assessment criteria related to environmental, economic, and societal sustainability, and discussed them in a focus group. This led to the highlighting of three native feedstocks – cellulose, lignin, and seaweed – that could be utilised in the UK to advance biobased economic transition, with scope for differing functionalities and the potential for multifaceted applications. We undertook detailed analysis of these three target technologies, using a framework that examined dimensions of functionality, sustainability, development, market and customers, scaling, industry (including investment and partnerships), and governance. While each technology exhibited variations by these dimensions, we found overall that current capabilities were constrained. At the same time, we also identified pathways towards sustainable development and scale up through circular economy models, stakeholder engagement, and changes in policies and economic incentives. Moreover, although Government has set out visions and targets, more specific attention to implementation. Coordination and funding are needed to support biobased production progression along circular value chains comprising feedstocks, biomanufacturing, marketable goods, and sustainable consumption, to foster innovation scale-up, support SMEs and de-risk technological development investment, along with longer-term funding to support systematic change. There is a need for better infrastructure for the accumulation and processing of waste for its use as feedstock and for legislative and policy changes to generate frameworks and incentives for sustainable biobased transition.

In the UK, resource availability and type of feedstock depends on the regional landscape and this needs to be considered in the development of a bioeconomy. For example, while Scotland has the highest availability of woodland and forest residues, Southeast England produces the highest fraction of municipal green waste. Feedstock availability, and the academic, technological, and industrial expertise in each area of the UK needs to be assessed. From this, a roadmap for implementation of a suitable and bespoke biobased strategy for each area can be developed. Ideally, this should be associated with devolution so that local stakeholders input into the management of local and regional resources, allowing sustainable customisation to address local biobased production demands and opportunities.

Any new regulations, policies, or incentives need new types of conversation, deliberation, and governance where there is greater interaction and transparency between policy makers, government, industry and publics. The post-pandemic environment has the potential to act as an unprecedented wake-up call to rectify previous failings. By utilising momentum from this crisis, where the status quo has been irreparably disrupted, bold governance and policy could leverage a systemic shift to integration and implementation of a sustainable biobased economy into the ‘new normal’.

We acknowledge limitations in this study. The targeted assessment approach, while informed by bibliometrics and analyses of available secondary data, depends on judgements made by the research team and selected experts and stakeholders about fit with key assessment criteria to select the most promising bioproduction technology approaches. The aim of the approach was not to exhaustively identify every possible technology, but rather to develop sufficiently diverse sets to inform discussion about the technologies in the context of broader economic, societal, and environmental sustainability and to highlight scaling challenges and policy issues. The use of a focus group to test and elaborate on these selections provides external review and input. In future work, the approach, with its focus on three streams of biobased production, could be scaled up and applied to a larger set of potential biobased technologies. However, going beyond a pilot study would require engaging a wider set of stakeholders to participate in assessments and multiple focus groups. Additionally, the study is focused on the UK. The findings do not necessarily apply to other national systems, although the approach used could be adapted for use elsewhere.

## Acknowledgements

CH and PS acknowledge support from the Biotechnology and Biological Sciences Research Council, UK [Grant Number BB/M017702/1] (Manchester Synthetic Biology Research Centre for Fine and Speciality Chemicals). PS further acknowledges support from the Engineering and Physical Sciences Research Council [Grant Number EP/S01778X/1] (Future Biomanufacturing Research Hub).

## Supplementary Materials (S)

### S1.1 Overview

This supplementary materials document provides additional details on the targeted assessment process.

### S1.2 Bioeconomy technologies: long list generation

An initial list of technologies that could contribute to bioeconomy development was formulated. Generating this long list was aided by a search for publications in the Web of Science (WoS) Core Collection. The search was targeted to the most recent period (2015-2020) due to the focus of the project on new and emerging technologies. Search terms included “Biobased”, “Bio-based”, “Biomaterial” or “Synthetic Biology”. Results were categorised into 25 WoS subject categories including Biotechnology Applied Microbiology, Biochemistry Molecular Biology, and Plant Science but excluding categories relating to medicine or medical applications that were outside of the scope of this work. Publications were then sorted within each of the remaining disciplines and ranked by citations. Manual review was undertaken of 10 high-ranking but diverse papers from each subject category. This was intended both to allow sufficient depth within a given subject category but to also allow breadth across disciplines. Following this, Google Scholar was used, alongside online patent, publication, and media resources, to identify other non-medical bioeconomy-relevant technologies.

The aim of this approach was not to exhaustively identify every possible technology, but rather to develop a sample long list that was sufficiently diverse, from different disciplines and feedstocks, to encompass a relatively broad section of the landscape of potential bioeconomy technologies. Concise profiles of potential technologies were developed by the author research team based on review of descriptions of the technology under development and available documentation (through publications, patents, or websites). There was a focus on technologies that were generic (i.e., not highly company specific) and either entering or relatively close to market – i.e., which could be implemented within, as a maximum, the next 5-10 years. A total of 50 potential bioeconomy technologies emerged (see Table S1).

### S1.3 Biobased production technologies: short-list generation

Technologies from the long list were further assessed on their ability to fulfil the following requirements: (1) Bio-based production technologies using the definition of “biobased” in the scope of the project (Non-food products derived from renewable biomass); (2) Nationally (UK) applicable – utilisation of native feedstock, resources, and expertise; (3) Economically beneficial – either by reducing dependence on imports, bolstering a particular market, or providing new goods or functionalities; (4) Socially positive – allowing local management of resources, job creation or other societal gains; and (5) Environmentally benign (or the potential to be so) – considering greenhouse gas (GHG) emissions toxicity, resource depletion, habitat destruction, water use, etc. In addition to author review, the list in progress was presented to experts in biotechnology, engineering biology, and science and technology policy (via workshops or seminars at The Future Biomanufacturing Research Hub; the Manchester Institute of Innovation Research; and the Manchester Synthetic Biology Research Center, and with members of the Institute for Food and Resource Economics at the University of Bonn). Of the original long list, 18 technologies were determined to fulfil the requirements criteria established and were included on the short-list of promising biobased production technologies. (Table S1).

**Table S1:**
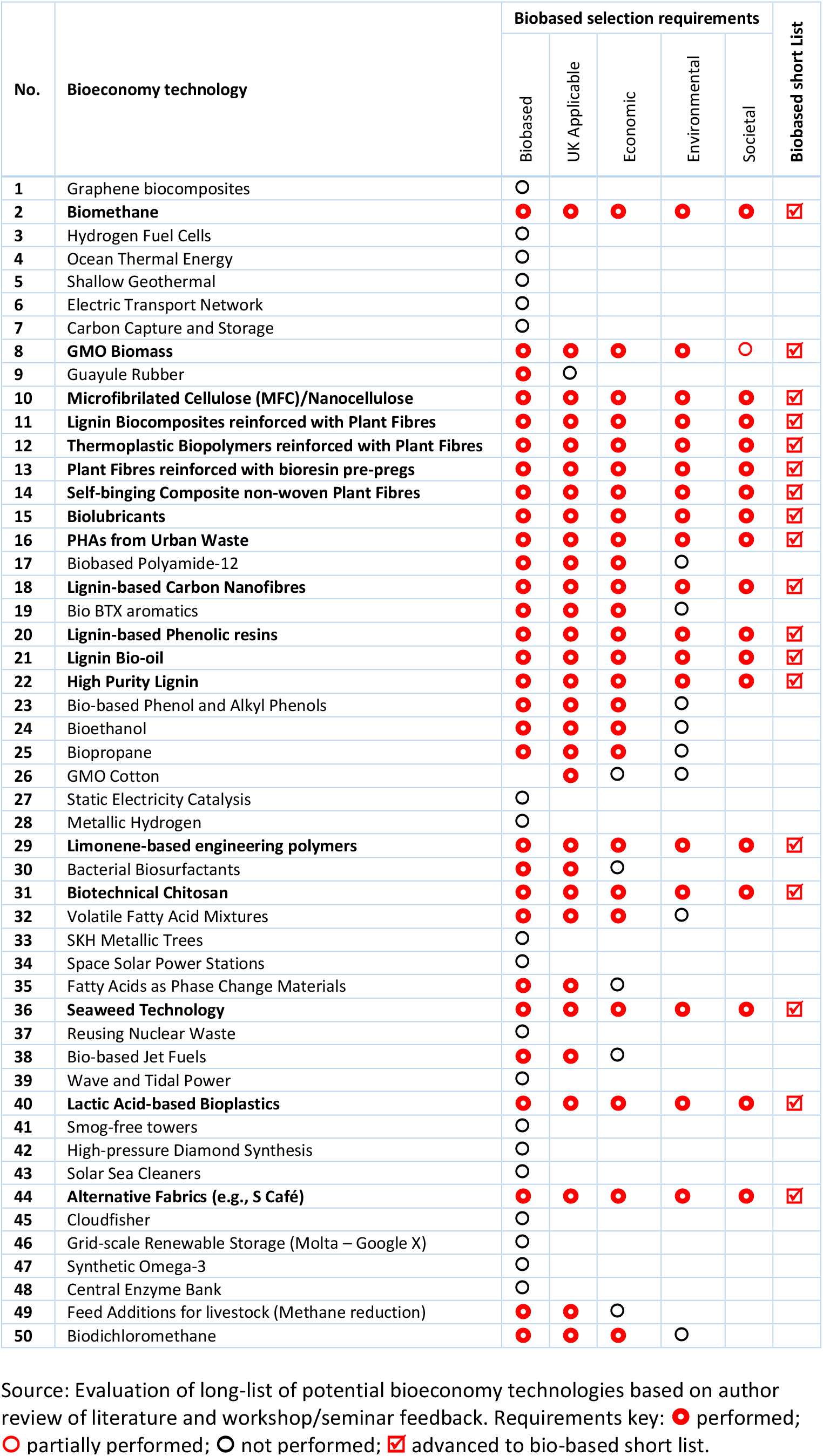
Long list bioeconomy technologies and bio-based production requirements fulfilment.

### S1.4 Focus Group

For the focus group, six technologies were selected from the short list as ‘exemplary technologies’ that represented a range of different feedstocks, low- and high-value products, and a diverse range of primary end uses. The technologies were:

1. Biomethane
2. High Purity Lignin
3. Microfibrilated Cellulose (MFC)/Nanocellulose
4. PHAs from fatty acids and urban waste
5. Seaweed technology
6. Bioethanol

Stakeholders represented industry leaders, academics, science platforms, think-tanks, government, policy, and feedstock providers, comprising 10 attendees plus two research team members. The session was hosted via Zoom in January 2021. Following an introduction of the project and explanation of its aims and selection criteria for bio-based technologies, focus group participants were split into two groups, each focussing on three of the exemplary technologies (Group A: Biomethane, MFC/nanocellulose, PHAs from urban waste; Group B: Bioethanol, High Purity Lignin, Seaweed Technology), and asked to answer three questions:

1. What are the specific challenges you see relevant to each of these specific technologies? E.g., technology/ management/ regulation, etc.
2. What are the main governance/ funding/ regulatory roadblocks you consider to be the most important and how can these be overcome?
3. Are there any other technologies you think should be included as exemplary case studies based on the UK’s native feedstocks/resources/expertise or policy drives?

Following discussion between stakeholders within the breakout groups, each group was asked to present their conclusions to the entire focus group, allowing further discussion of all technologies. After the focus group, feedback was provided to all participants, allowing them to provide additional insight or resources, and generate further discussion.

### S1.5 Case Studies

Case studies were created for lignin, cellulose, and seaweed using online resources, peer-reviewed publications, patents, and direct communication with field experts and industry. The case studies focussed on the following probes (Figure S1):

1. Functionality: Technical specification, novel functions, and comparison to the status quo.
2. Sustainability: Source of feedstock, ability of UK to provide / import, and sustainability/environmental issues of production and consumption.
3. Development: Current use, and alternative and developing methods of utilisation, extraction, and valorisation.
4. Market and Customers: Affected markets and customers, current and potential scale / growth / locations of the market, and potential market competitors.
5. Scaling: Potential for and requirements of scalability of the extraction/ valorisation process, and examples of technologies in development.
6. Industry/ Investment/ Partners: Current industry structure, potential for redistributed (small-scale, decentralised) production, and capital requirements.
7. Governance: Regulatory aspects, public acceptance and possible opposition to this feedstock / technology.

**Figure S1:**
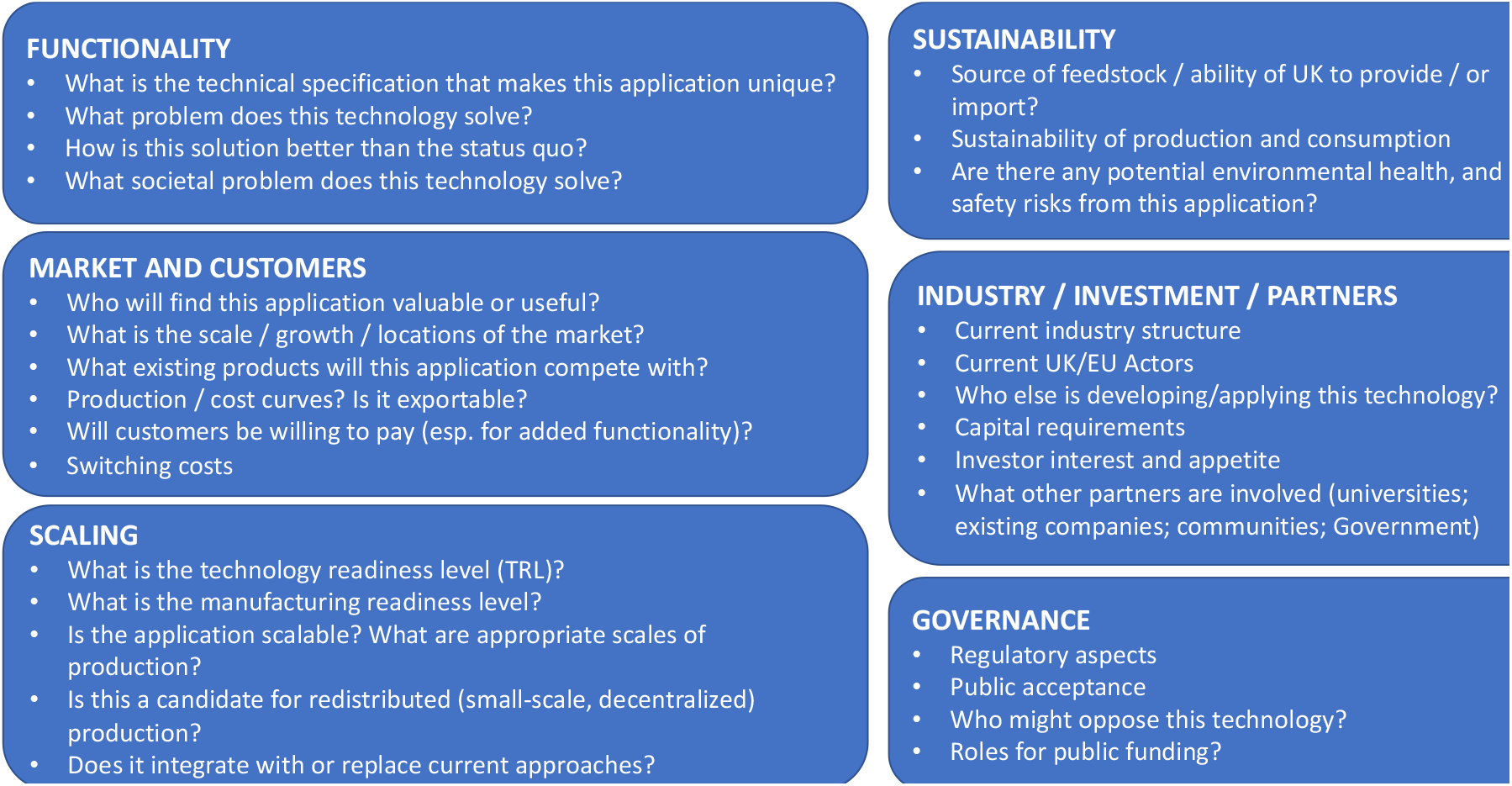
Case study probes.

## References

AB-SIG, 2013. A UK roadmap for algal technologies. Collated for the NERC-TSB Algal Bioenergy Special Interest Group by B. Schlarb-Ridley and B. Parker, Adapt, pp. 75.

Becker, J., Wittmann, C., 2019. A field of dreams: Lignin valorization into chemicals, materials, fuels, and health-care products. Biotech. Advances. doi: 10.1016/j.biotechadv.2019.02.016

BEIS, 2018. Bioeconomy strategy: 2018 to 2030. Department for Business, Energy & Industrial Strategy. https://www.gov.uk/government/publications/bioeconomy-strategy-2018-to-2030.

BEIS, 2021. UK Innovation Strategy: leading the future by creating it. Department for Business, Energy & Industrial Strategy. https://www.gov.uk/government/publications/uk-innovation-strategy-leading-the-future-by-creating-it.

Boehlje, M., Bröring, S., 2011 The increasing multifunctionality of agricultural raw materials: Three dilemmas for innovation and adoption, Int. Food Agribus. Man., 14: 1–16

Boerjan W, Ralph J, Baucher M., 2003. Lignin biosynthesis. Annu Rev Plant Biol., 54: 519–546

Boudet, A.M., Kajita, S., Grima-Pettenati, J., Goffner, D., 2003. Lignins and lignocellulosics: a better control of synthesis for new and improved uses. Trends in Plant Science, 8: 576–581

Bringezu, S., O’Brien, M., Schütz, H., 2012. Beyond biofuels: Assessing global land use for domestic consumption of biomass. A conceptual and empirical contribution to sustainable management of global resources. Land Use Policy, 29, 224–232

Buschke, N., Schröder, H., Wittmann, C., 2011. Metabolic engineering of Corynebacteriym glutamicum for production of 1,5-diaminopentane from hemicellulose, Biotech. J., 6, 306–317

Buschmann, A.H., Camus, C., Infante, J. et al., 2017. Seaweed production: an overview of the global state of exploitation, farming and emerging research activity. European Journal of Phycology. 52: 391–406

Cabral, P., Levrel, H., Viard, F., Frangoudes, K., Girard, S., Scemama, P., 2016. Ecosystem services assessment and compensation costs for installing seaweed farms. Marine Policy, 71: 157–165.

Cao, L., Yu, I.K.M, Liu, Y., et al., 2018. Lignin valorization for the production of renewable chemicals: State-of-the-art review and future prospects. Bioresour. Technol. 269: 465–475

Cefas, 2016. Seaweed in the UK and abroad – status, products, limitations, gas and Cefas role. Centre for Environment, Fisheries and Aquaculture Science. https://www.gov.uk/government/publications/the-seaweed-industry-in-the-uk-and-abroad

Chemical & Engineering News Archive, 1984. Lignin Is Coming of Age for Use in Polymeric Materials, Chem. Eng. News. 62: 19–20

Chen, Z., Wan, C., 2018. Ultrafast fractionation of lignocellulosic biomass by microwave-assisted deep eutectic solvent pretreatment. Bioresour Technol. 250: 532–537

Clark, J.H., Deswarte, F., 2008. The Biorefinery Concept – An Integrated Approach. John Wiley & Sons, Chichester, UK.

Conrad, J.M., Clark, C.W., 1987. Natural Resource Economics: Notes and Problems. Cambridge University Press, Cambridge, UK

D’Amato, D., Veijonaho, S., Toppinen, A., 2020. Towards sustainability? Forest-based circular bioeconomy business models in Finnish SMEs, Forest Policy and Economics, 127: 102455 doi:10.1016/j.forpol.2018.12.004

De Besi, M., McCormick, K., 2015. Towards a bioeconomy in Europe. National, regional and industrial strategies. Sustainability, 7, 10461–10478

De Wild, P., Huijgen, W.J.J., Gosselink, R.J.A., 2014. Lignin pyrolysis for profitable lignocellulosic biorefineries. Biofuels, Bioprod. And Bioref. DOI: 10.1002/bbb.1474

DEFRA, 2021. UK statistics on waste. Government Statistical Service. https://assets.publishing.service.gov.uk/government/uploads/system/uploads/attachment_data/file/1002246/UK_stats_on_waste_statistical_notice_July2021_accessible_FINAL.pdf

Duarte C.M., Wu J., Xiao X., Bruhn A., Krause-Jensen, D., 2017. Can Seaweed Farming Play a Role in Climate Change Mitigation and Adaption? Frontiers in Marine Science, 4:100, doi: 10.3389/fmars.2017.00100

Duarte, C. M., Losada, I. J., Hendriks, I. E., Mazarrasa, I., Marbà, N., 2013. The role of coastal plant communities for climate change mitigation and adaptation. Nat. Clim. Change, 3, 961–968. doi: 10.1038/nclimate1970

Duarte, C. M., Middelburg, J., Caraco, N., 2005. Major role of marine vegetation on the oceanic carbon cycle. Biogeosciences, 2, 1–8. doi: 10.5194/bg-2-1-2005

EC, 2012. Innovating for Sustainable Growth – A Bioeconomy for Europe, Research and Innovation, European Commission, Brussels.

EC, 2015. Sustainable Agriculture, Forestry and Fisheries in the Bioeconomy – A challenge for Europe, 4th SCAR Foresight Exercise. European Commission, Brussels.

EC, 2018. A Sustainable Bioeconomy for Europe: Strengthening the Connection between Economy. European Commission, Brussels.

Ek, M., Gellerstedt, G., Henriksson, G. (Eds), 2009. Pulping Chemistry and Technology Vol. 2, Walter de Gruyter, Berlin

Enriquez-Cabot, J., 1998. Genomics and the world’s economy. Science 281, 925–926

EP, 2009a. Directive 2009/28/EC of the European Parliament and of the Council of 23 April 2009 on the Promotion of the Use of Energy from Renewable Sources and Amending and Subsequently Repealing Directives 2001/77/EC and 2003/30/EC; European Parliament: Strasbourg.

EP, 2009b. Directive 2009/30/EC of the European Parliament and of the Council of 23 April 2009 amending Directive 98/70/EC as Regards the Specification of Petrol, Diesel and Gas-Oil and Introducing a Mechanism to Monitor and Reduce Greenhouse Gas Emissions and Amending Council Directive 1999/32/EC as Regards the Specification of Fuel Used by Inland Waterway Vessels and Repealing Directive 93/12/EEC; European Parliament: Strasbourg.

ERRMA, 2007. Accelerating the development of the market for bio-based products in Europe. Report of the Taskforce of Bio-Based Products. European Renewable Resources and Materials Association,

Essel, R., Carus, M., 2014. Increasing resource efficient by cascading use of biomass, Rural, 21: 28–29

FAO (2020) The State of World Fisheries and Aquaculture 2020. Sustainability in Action. Rome. 10.4060/ca9229en

FAO, 2016. How Sustainability Is Addressed in Official Bioeconomy Strategies at International, National and Regional Levels; Food and Agriculture Organization of the United Nations: Rome, Italy.

FBI, 2020. Cellulose Market Size, Share & Industry Analysis By Derivative Type (Commodity Cellulose Pulp, Cellulose Fibers, Cellulose Ethers, Microcrystalline Cellulose, Nanocellulose, and Others), By End-Use Industry (Textile, Food, Chemical Synthesis, Pharmaceuticals, Construction, Paper & Pulp, Paints & Coatings, and Others), Regional Forecast, 2019-2026. Report IB: FBI102062. Fortune Business Insights. https://www.fortunebusinessinsights.com/cellulose-market-102062

Ferdouse, F., Holdt, S.L., Smith, R., Murúa, P., Yang, Z., 2018. The global status of seaweed production, trade and utilization. GLOBEFISH Research Programme. Rome. Volume 124. https://www.proquest.com/openview/63a9872d1ea30c63f92d5d8acfcd6e35/1?pq-origsite=gscholar&cbl=237312

Fernandez-Rodriguez, J., Erdocia, X., Sanchez, C., Alriols, M.G., Labidi, J., 2017. Lignin depolymerization for phenolic monomers production by sustainable processes. Energy Chem. 26: 622–631

Forestry Commission, 2020. Provisional Woodland Statistics 2020 Edition, https://www.forestresearch.gov.uk/documents/7647/PWS_2020_ggXAYhv.pdf

Fourqurean, J. W., Duarte, C. M., Kennedy, H., Marbà, N., Holmer, M., Mateo, M. A., et al., 2012. Seagrass ecosystems as a globally significant carbon stock. Nat. Geosci., 5, 505–509. doi: 10.1038/ngeo1477

Fritsche, U.R., Irarte, L., 2014. Sustainability criteria and indicators for the bio-based economy in Europe: State of discussion and way forward, Energies, 7: 6825–6836

GMI, 2020. Cellulose Market Size, By Source (Natural, [Crops, Cotton, Jute, Hemp, Flax, Corn, Wheat], Fruits, [Apple, Peaches, Strawberries, Treewood [Hardwood, Softwood], Synthetic), By Modification (Unmodified, Modified, [Microcrystalline Cellulose, Cellulose Ethers, Oxidized), By Manufacturing Process (Viscose, [Acid Sulfite, Prehydrolysis Kraft], Cellulose Ethers), By Purity (Above 95%, 85% - 95%, Below 85%), By Application (Food, Pharmaceuticals, Paper, Cosmetics, Textiles), Industry Analysis Report, Regional Outlook, Application Growth Potential, Price Trends, Competitive Market Share & Forecast, 2020 – 2026. Global Market Insights. https://www.gminsights.com/industry-analysis/cellulose-market

Gomez-Zavaglia, A., Prieto Lage, M.A., Jimenez-Lopez, C., Mejuto, J.C., Simal-Gandara, J., 2019. The Potential of Seaweeds as a Source of Functional Ingredients of Prebiotic and Antioxidant Value, Antioxidants, 8: 406

Hepworth, D.G., Bruce, D.M., 2000. A method of calculating the mechanical properties of nanoscopic plant cell wall components from tissue properties. J. of Materials Science, 23: 5861–5865

HM Treasury, 2021. Build Back Better: our plan for growth. CP 401. 3 March.

Holland, C., Perzon, A., Cassland, P.R.C., et al., 2019. Nanofibers Produced from Agro-Industrial Plant Waste Using Entirely Enzymatic Pretreatments. Biomacromolecules. 20: 443–453

IPCC, 2018. Special Report: Global Warming of 1.5 □ C. Intergovernmental Panel on Climate Change https://www.ipcc.ch/sr15/

Keep, M., 2022. Economic update: Economy was back to pre-pandemic level before Omicron. House of Commons Library, 25 January. https://commonslibrary.parliament.uk/economic-update-economy-was-back-to-pre-pandemic-level-before-omicron/

Kershaw E.H., Hartley S., McLeod C., Polson P., 2020. The Sustainable Path to a Circular Economy, Trends in Biotechnology, 39(6):542–545. doi: 10.1016/j.tibtech.2020.10.015.

Khan, A., Naire, V., Colmenares, J.C., Gläsier, R., 2018. Lignin-based composite materials for photocatalysis and photovoltaics. Rop. Curr. Chem. 376: 20

Kircher, M., 2012. The transition to a bio-economy: National perspectives. Biofuel. Bioprod. Biorefin. 6, 240–245.

Kitchen L., Marsden T., 2011. Constructing sustainable communities: a theoretical exploration of the bio-economy and eco-economy paradigms, Local Environment, 16: 753–769

Klemm, D., Heublein, B., Fink, H-P., Bohn, A., 2005. Cellulose: Fascinating Biopolymer and Sustainable Raw Material. 44: 3358–3393

Kohlstedt, M., Starck, S., Barton, N., et al., 2018. From lignin to nylon: Cascaded chemical and biochemical conversion using metabolically engineered Pseudomonas putida. Metab. Eng. 47: 279–293

Landeweerd, L., Surette, M., van Driel, C., 2011. From petrochemistry to biotech: A European perspective on the bio-based economy. Interface Focus, 1, 189–195

Langeveld, J.W.A., Dixon, J., Jaworski, J.F., 2010. Development perspectives of the biobased economy: A review. Crop Sci., 50, S-142–S-151

Levidow, L., Birch, K., Papaioannou, T., 2012. EU agri-innovation policy: two contending visions of the bio-economy. Crit. Pol. Stud. 6: 40–65. DOI: 10.1080/19460171.2012.659881

Li, X., Zhang, Z., Qu, S., Liang, G., Zhao, N., Sun, J., et al., 2016. Breeding of an intraspecific kelp hybrid Dongfang no. 6 (Saccharina japonica, Phaeophyceae, Laminariales) for suitable processing products and evaluation of its culture performance. Journal of Applied Phycology 28: 439–447.

Liu, X., Ma, Y., Zhang, M., 2015. Research advances in expansins and expansion-like proteins involved in lignocellulose degradation. Biotechnol. Lett. 37: 1541

Losacker, S., Heiden, S., Liefner I., Lucas, H. 2023. Rethinking bioeconomy innovation in sustainability transitions. Technology in Society 74, 102291. doi: 10.1016/j.techsoc.2023.102291.

Makkar, H. P., Tran, G., Heuzé, V., Giger-Reverdin, S., Lessire, M., Lebas, F., et al., 2016. Seaweeds for livestock diets: a review. Anim. Feed Sci. Technol. 212, 1–17. doi: 10.1016/j.anifeedsci.2015.09.018

Martín-Blanco, C., Zamorano, M., Lizárraga, C., Molina-Moreno, V., 2022. The Impact of COVID-19 on the Sustainable Development Goals: Achievements and Expectations. Int J Environ Res Public Health 19(23):16266. doi: 10.3390/ijerph192316266.

Matthews, N.E., Stamford, L., Shapira, P., 2019. Aligning sustainability assessment with responsible research and innovation: Towards a framework for Constructive Sustainability Assessment. Sustainable Production and Consumption, 20, 58–73, doi: 10.1016/j.spc.2019.05.002.

Mazarrasa, I., Olsen, Y. S., Mayol, E., Marbà, N., Duarte, C. M., 2014. Global unbalance in seaweed production, research effort and biotechnology markets. Biotechnol. Adv. 32, 1028–1036. doi: 10.1016/j.biotechadv.2014.05.002

McCollum, D.L., Zhou, W., Bertram, C., Bosetti, V., et al., 2018. Energy investment needs for fulfilling the Paris Agreement and achieving the Sustainable Development Goals. Nature Energy, 3, 589–599

McLeod, E., Chmura, G. L., Bouillon, S., Salm, R., Björk, M., Duarte, C. M., et al., 2011. A blueprint for blue carbon: towards an improved understanding of the role of vegetated coastal habitats in sequestering CO2. Front. Ecol. Environ., 9, 552–560. doi: 10.1890/110004

MHCL, 2020. Land Use in England, 2018. Planning. Ministry of Housing, Communities & Local Government, official statistics release.

Michels, J., Wagemann, K., 2010. The German Lignocellulose Feedstock Biorefinery Project. Biofuels Bioprod. Biorefin. 4: 263–267

Milone, P., Ventura, F., 2010. Networking the rural: the future of green regions in Europe, The Netherlands: Van Gorcum.

Mouritsen, O. G. 2013. Seaweeds edible, available and sustainable. The University of Chicago Press, Chicago and London; pp. 287

Naidoo, R., Fisher, B., 2020. Reset Sustainable Development Goals for a pandemic world. Nature. 583: 198–201

Naseem, A., Rabasum, S., Zia, K.M., Zuber, M., Ali, M., Noreen, A., 2016. Lignin-derivatives based polymers, blends and composites: A review. Int. J. Biol. Macromol., 93: 296–313

Nizami, A.S., Mohanakrishna, G., Mishra, U., Pant, D., 2016. Trends and SustainabilityCriteria for the Liquid Biofuels. Biofuels: Production and Future Perspectives. CRCPress, pp. 59–95. 10.1201/9781315370743-1

Nizami A.S., Rehan M., Waqas M., Naqvi M., Ouda O.K.M., Shahzad K., Miandad R., Khan M.Z., Syamsiro M., Ismail I.M.I. & Pant D., 2017a. Waste biorefineries: Enabling circular economies in developing countries, Bioresource Technology, 241: 1101–1117

Nizami, A.S., Shazad, K.m Rehan, M., Ouda, O.K.M., Khan, M.Z., Ismail, I.M.I., Almeelbi, T., Basahi, J.M., Demirbas, A. (2017b) Developing waste biorefinery in Makkah: a way forward to convert urban waste into renewable energy. Appl. Energy. 186: 189–196

NFCC (2014) Lignocellulosic Feedstock in the UK. York, UK. https://www.nnfcc.co.uk/files/mydocs/LBNet%20Lignocellulosic%20feedstockin%20the%20UK_Nov%202014.pdf

Nuss, P., Gardner, K.H., 2013. Attributional life cycle assessment (ALCA) of polyitaconic acid production from northeast US softwood biomass. Int. J. Life Cycle Ass., 18, 603–612.

O’Doherty, J. V., Dillon, S., Figat, S., Callan, J. J., Sweeney, T., 2010. The effects of lactose inclusion and seaweed extract derived from Laminaria spp. on performance, digestibility of diet components and microbial populations in newly weaned pigs. Anim. Feed Sci. Technol., 157, 173–180. doi: 10.1016/j.anifeedsci.2010.03.004

OECD, 2020. Building Back Better: A Sustainable, Resilient Recovery after COVID-19. Organisation for Economic Cooperation and Development, Paris.

OECD, 2023. Net Zero+: Climate and Economic Resilience in a Changing World. Organisation for Economic Cooperation and Development, Paris. doi: 10.1787/da477dda-en.

Oldekop, J.A., Horner, R., Hulme, D., et al. (2020) COVID-19 and the case for global development. World Dev. 134: 105044. doi: 10.1016/j.worlddev.2020.105044

ONS, 2020. GDP first quarterly estimate, UK: April to June. Office for National Statistics. 2020https://www.ons.gov.uk/economy/grossdomesticproductgdp/bulletins/gdpfirstquarterlyestimateuk/apriltojune2020

O’Shea, C. J., McAlpine, P., Sweeney, T., Varley, P. F., O’Doherty, J. V., 2014. Effect of the interaction of seaweed extracts containing laminarin and fucoidan with zinc oxide on the growth performance, digestibility and faecal characteristics of growing piglets. Br. J. Nutr., 111, 798–807. doi: 10.1017/S0007114513003280

Overhage, J., Steinbüchel, A., Priefert, H., 2003. Highly efficient biotransformation of eugenol to ferulic acid and further conversion to vanillin in recombinant strains of Escherichia coli. Appl. Environ. Microbiol. 69: 6569–6576

Pedroli, B., Elbersen, B., Frederiksen, P., Grandin, U., Heikkilä, R., Krogh, P.H., Izakovicova, Z., Johansen, A., Meiresonne, L., Spijker, J., 2013. Is energy cropping in Europe compatible with biodiversity? Opportunities and threats to biodiversity from land-based production of biomass for bioenergy purposes. Biomass Bioenergy, 55: 73–86

Pfau S.F., Hagens J.E., Dankbaar B. Smits A.J.M., 2014, Visions of Sustainability in the Bioeconomy Research. Sustainability, 6, 1222–1249

Philp, J., 2018. The bioeconomy, the challenge of the century for policy makers. N. Biotechnol. 40: 11–19.

Philp, J., Winickoff, D.E., 2018. Realising the Circular Bioeconomy. OECD Science, Technology and Industry Policy Papers, No. 60

Popescu, I., 2014. Industrial biotechnology in the European union: identifying the best pathways to boost growth of the bioeconomy. Ind. Biotechnol, 10, 376–378. doi: 10.1089/ind.2014.1537

Priefer C., Jörissen J., Frör O., 2017. Pathways to Shape the Bioeconomy, Resources, 6, doi: 10.3390/resources6010010

Pye, E.K., 2010. Industrial lignin production and application. In: Kamm, B., Gruber, P.R., Kamm, M. (Eds). Biorefineries – Industrial processes and products. Wiley-VCH Verlag GmbH & Co. KGaA, Weinheim. pp 165–200

Raghu, S., Spencer, J.L., Davis, A.S., Wiedenmann, R.N., 2011. Ecological consideration in the sustainable development of terrestrial biofuel crops. Curr. Opin. Environ. Sustain., 3, 15–23.

Ragnar, M., Henriksson, G., Lindström, M.E., Wimby, M., Blechschmidt, J., Heinemann, S., 2014. Pulp. In: Ullmann’s Encyclopedia of Industrial Chemistry, Wiley-VCH, Weinheim

Ralph, J., 2010. Hydroxycinnamates in lignification. Phytochem. Rev., 9, 65–83

Rey-Crespo, F., Lopez-Alonso, M., Miranda, M., 2014. The use of seaweed from the Galician coast as a mineral supplement in organic dairy cattle. Animal 8, 580–586. doi: 10.1017/s1751731113002474

Rinaldi, R., Jastrzebski, R., Clough, M.T., Ralph, J., Kennema, M., Bruijnincx, P.C., Weckhuysen, B.M., 2016. Paving the way for lignin valorisation: recent advances in bioengineering, biorefining and catalysis. Angew. Chem, Int. Ed. Engl., 55, 8164–8215

Roberts, D. A., Paul, N. A., Dworjanyn, S. A., Bird, M. I., de Nys, R., 2015. Biochar from commercially cultivated seaweed for soil amelioration. Sci. Rep. 5:9665. doi: 10.1038/srep09665

Robertson, G. P., Paul, E. A., Harwood, R.R., 2000. Greenhouse gases in intensive agriculture: contributions of individual gases to the radiative forcing of the atmosphere. Science 289, 922–1925. doi: 10.1126/science.289.5486.1922

Robinson, N., Winberg, P. Kirkendale, L., 2013. Genetic improvement of macroalgae: status to date and needs for the future. Journal of Applied Phycology, 25: 703–716.

Ronzon, T., Iost, S. Philippidis, G., 2022. Has the European Union entered a bioeconomy transition? Combining an output-based approach with a shift-share analysis. Environment, Development, and Sustainability. 10.1007/s10668-021-01780-8

Rosas, J.M., Berenguer, R., Valero-Romero, M.J., Rodríguez-Mirasol, J., Cordero, T., 2014. Preparation of different carbon materials by thermochemical conversion of lignin. Front. Mater. 10.3389/fmats.2014.00029

Rosengrant, M.W., Ringler, C., Zhu, T., Tokgoz, S., Bhandary, P. (2013) Water and food in the bioeconomy: Challenges and opportunities for development. Agric. Econ. 44, 139–150.

Schutyser, W., Renders, T., van den Bosch, S., et al., 2018. Chemicals from lignin: an interplay of lignocellulose fractionation, depolymerisation, and upgrading. Chem. Soc. Rev. 47, 852–908

SDC, 2011. Looking Back, Looking Forward—Sustainability and UK Food Policy 2000–2011. Sustainable Development Commission: London.

Searle, S., Mallins, C., 2013. Availability of celluloisic residues and wastes in the EU, White Paper. The International Council on Clean Transportation, Washington, DC.

Sharholy, M., Ahmad, K., Mahmood, G., Trivedi, R.C., 2008. Municipal solid waste management in Indian cities, a review. Waste Management. 28, 459–467

Sheppard, A.W., Raghu, S., Begley, C., Genovesi, P., de Barro, P., Tasker, A., Roberts, B., 2011. Biosecurity as an integral part of the new bioeconomy: A path to a more sustainable future. Curr. Opin. Environ. Sustain., 3, 105–111.

Sims, R., Mabee, W., Saddler, J., Taylor., M., 2010. An overview of second generation biofuel technologies. Bioresour. Technol., 101, 1570–1580

Smale, D. A., Burrows, M. T., Moore, P., O’Connor, N., and Hawkins, S. J., 2013. Threats and knowledge gaps for ecosystem services provided by kelp forests: a northeast Atlantic perspective. Ecol. Evol. 3, 4016–4038

Smith, B. E., 2002. Nitrogenase reveals its inner secrets. Science 297, 1654–1655. doi: 10.1126/science.1076659

Sonnino, R., Marsden, T.K., 2006. Beyond the divide: rethinking relationships between alternative and conventional food networks in Europe. Journal of Economic Geography, 6, 2, 181–199.

Sonoki, T., Morooka, M., Sakamoto, K., et al., 2014. Enhancement of protocatechuate decarboxylase activity for the effective production of muconate from lignin-related aromatic compounds. J. Biotechnol. 192, 71–77

Stark, S., Biber-Freudenberger, L., Dietz, T., Escobar, N., Förster, J.J., Henderson, J., Laibach, N., Börner, J. 2022. Sustainability implications of transformation pathways for the bioeconomy. Sustainable Production and Consumption 29, 215–227. doi: 10.1016/j.spc.2021.10.011

Stegmann, P., Londo, M., Junginger, M., 2020. The circular bioeconomy: Its elements and role in European bioeconomy clusters, Resources, Conservation & Recycling, 10.1016/j.rcrx.2019.100029

Steneck, R. S., Graham, M. H., Bourq, B. J., Corbett, D., Erlandson, J. M., 2002. Kelp forest ecosystems, biodiversity, stability resilience and future. Environ. Conserv., 29, 436–459

Strassberger, Z., Tanase, S., Rothenberg, G., 2014. The pros and cons of lignin valorisation in an integrated biorefinery. RSC Adv. 4: 25310–25318.

Thielemans, W., Can, E., Morye, S.S., Wool, R.P., 2002. Novel applications of lignin in composite materials, J. of Applied Polymer Sci., 83, 323–331

UK Parliament (2008) Climate Change Act 2008 (c27). https://www.legislation.gov.uk/ukpga/2008/27/contents

van den Bosch, S., Koelewijn, S.F., Renders, T., Van den Bossche, G. Vangeel, T., Schutyser, W., Sels, B.F. 2018 Catalytic Strategies Towards Lignin-Derived Chemicals. Top. Curr. Chem., 376: 36

Vardon, D.R., Franden, M.A., Johnson, C.W., et al., 2015. Adipic acid production from lignin. Energy Environ. Sci. 8, 617–628

Vásquez, J.A.J., Zuñiga, S., Tala, F., Piaget, N., Rodríguez, D.C. & Vega, J.M.A., 2014. Economic valuation of kelp forests in northern Chile: values of goods and services of the ecosystem. Journal of Applied Phycology, 26: 1081–1088

Venkata Mohan S., Nikhil G.N., Chiranjeevi P., Nagendranatha Reddy C., Rohit M.V., Naresh Kumar A., Sakar O., 2016. Waste biorefinery models towards sustainable circular bioeconomy: Critical review and future perspectives, Bioresource Technology, 215, 2–12

Viell, J., Harwardt, A., Seiler, J., Marquardt, W. (2013) Is biomass fractionation by Organosolv-like processes economically viable? A conceptual design study, Bioresour. Technol. 150: 89–97.

Vohra, K., Vodonos, A., Schwartz, J., Marais, E.A, Sulprizio, M.P., Mickley, L.J., 2021. Global mortality from outdoor fine particulate pollution generated by fossil fuel combustion: Results from GEOS-Chem, Environmental Research, 195, 110754

West, J., Calumpong, H. P., Martin, G., 2016. Seaweeds. Chapter 14 in The First Global Integrated Marine Assessment. World Ocean Assessment I. United Nations. Available at: http://www.un.org/Depts/los/global_reporting/WOA_RegProcess.htm.

WRAP (2019) Food Waste Reduction Roadmap progress report 2019. https://wrap.org.uk/sites/default/files/2020-07/WRAP-Food-Waste-Reduction_Roadmap_Progress-Report-2019.pdf

Wu, W., Dutta, T., Varman, A.M., et al., 2017. Lignin Valorization: Two Hybrid Biochemical Routes for the Conversion of Polymeric Lignin into Value-added Chemicals. Nature Scientific Reports. 7, 8420

Zhang, Q.S., Tang, X.X., Cong, Y.Z., Qu, S.C., Luo, S.J., Yang, G.P., 2007. Breeding of an elite Laminaria variety 90-1 through inter-specific gametophyte crossing. Journal of Applied Phycology, 19, 303–311.

Zilberman D., Kim E., Kirschner S., Kaplan S., Reeves J., 2013. Technology and the future bioeconomy, Agricultural Economics, 44, 95–102.

